# Preparation duration shapes the goal-directed tuning of stretch reflex responses

**DOI:** 10.1101/2025.04.29.651111

**Authors:** Robin Rohlén, Frida Torell, Michael Dimitriou

**Author notes:** Correspondence to Robin Rohlén. **Statements and Declarations:** The authors have no competing interests to declare that are relevant to the content of this study. **Author contributions:** All authors contributed to the study’s conception and design. RR performed data curation and analysis. RR wrote the first draft of the manuscript. All authors reviewed and edited the manuscript. All authors read and approved the final manuscript.

## Abstract

Stretch reflex responses are important for counteracting perturbations, and the modulation of stretch reflex gains can assist the voluntary movement quality and performance. Recent findings suggest that movement preparation includes goal-directed tuning of muscle spindles, facilitating a concurring modulation of reflex gains. In the context of delayed-reach, goal-directed modulation of stretch reflex gains is present when the homonymous muscle is unloaded (i.e., antagonist load) and the preparatory delay is sufficiently long (>250 ms and <750 ms). This study examined how multiple preparatory delays impact the goal-directed modulation of short- and long-latency stretch reflex (SLR and LLR) gains of the pectoralis, anterior and posterior deltoid muscles in a delayed-reach task. We used electromyographic signals to quantify stretch reflex responses of healthy subjects that experienced haptic perturbations induced by a robotic manipulandum in a delayed-reach task. We found that a preparatory delay of 300 ms is sufficient for goal-directed tuning of SLR responses in the pectoralis and posterior deltoid muscles, and 400 ms allows for even stronger goal-directed tuning of stretch reflex responses. In contrast, there was no goal-directed modulation in the SLR of the anterior deltoid. Tuning of LLR gains was robustly goal-dependent throughout, with negligible modulation across preparatory delays. Our results clarify the minimum preparatory time required for goal-directed tuning of stretch reflex gains and characterise the relationship between reflex gains and preparation duration. This relationship is not strictly linear, likely reflecting the interplay of multiple feedback mechanisms functioning at different time frames.

## Introduction

Voluntary movements are typically preceded by a preparatory phase in which neural signals are modulated in anticipation of motor execution (Kutas and Donchin, 1974; Wise, 1985; Ghez *et al*., 1997). During this preparatory period, neural activity neural activity has been shown to predict several movement parameters, such as direction (Tanji and Evarts, 1976), velocity (Churchland, Santhanam and Shenoy, 2006), extent of reach (Messier and Kalaska, 2000), trajectory (Hocherman and Wise, 1991), and the spatial location of the visual target (Batista *et al*., 2007). This phase of preparation also includes the formulation of anticipatory motor commands and the tuning of feedback systems that influence long-latency stretch reflex responses (LLRs) before any physical action begins (Todorov and Jordan, 2002; Shadmehr and Krakauer, 2008; Wagner and Smith, 2008; Ahmadi-Pajouh, Towhidkhah and Shadmehr, 2012; Yeo, Franklin and Wolpert, 2016).

When an unexpected stretch occurs in the upper limb muscles, it can elicit a short-latency reflex (SLR) within approximately 25 ms and a subsequent LLR around 50 ms post-perturbation (Hammond, 1956; Marsden, Merton and Morton, 1972). Both reflexes are classified as involuntary, with volitional control of movement emerging around 100 ms after the initial sensory input (Yang *et al*., 2011). Historically, these stretch reflexes have been understood as mechanisms that contribute to the maintenance of posture by counteracting unanticipated disturbances (Nichols and Houk, 1976). However, it is now recognised that reflex responses can also be automatically modulated in a goal-dependent manner during voluntary actions. For example, reflex modulation has been observed in response to variations in target features (Nashed, Crevecoeur and Scott, 2012), the presence of obstacles (Nashed, Crevecoeur and Scott, 2014), and dynamic or static target properties (Cluff and Scott, 2015), as well as based on directional cues (Pruszynski, Kurtzer and Scott, 2008; Scott, 2016).

In the context of upper limb control, the SLR typically exhibits a dependency on initial muscle load – an effect known as automatic gain scaling – while the LLR demonstrates more nuanced task-specific tuning (Pruszynski *et al*., 2009, 2011). Recent evidence further differentiates the early and late components of the LLR: the early phase appears to include a stabilising element that operates independently of the voluntary action being prepared, while the later phase reflects task-specific adaptation (Lee and Perreault, 2019). These insights support the idea that components of the LLR can be flexibly modulated and may not be purely reflexive in nature (Shemmell, Krutky and Perreault, 2010). As previously described, feedback mechanisms responsible for modulating LLRs are believed to be configured during the preparatory stage of reaching (Ahmadi-Pajouh, Towhidkhah and Shadmehr, 2012). Although the precise mechanisms linking preparatory neural activity to optimised motor output remain elusive, current theories suggest that preparatory activity helps establish the initial neural state necessary for executing goal-directed movement (Churchland *et al*., 2010, 2012).

Recent studies suggest that tuning of muscle spindle sensitivity represent an independent component of movement preparation, contributing to the modulation of reflex gain in a task-relevant way (Papaioannou and Dimitriou, 2021). This view supports the involvement of two distinct pathways in preparatory control: one targeting alpha motor neurons and another regulating gamma motor neurons (Dimitriou, 2021, 2022). By modulating the responsiveness of spindles and associated reflexes via gamma control, the nervous system can influence sensorimotor feedback independently of the force generated during preparation. For instance, the alignment of reflex gain modulation with target direction facilitates more efficient movement execution (Torell *et al*., 2023). This tuning of stretch reflex gains – observed even at the SLR level – was most evident when the involved muscle was unloaded and when a preparatory delay of 750 ms was present, with 250 ms being too short. However, the minimum time required for such tuning to emerge and how this modulation develops over time remain open questions.

In this study, we examine how a series of multiple preparatory delays impact the goal-directed modulation of stretch reflex gains of the pectoralis, anterior and posterior deltoid muscles. To achieve this, we used electromyographic (EMG) signals to quantify stretch reflex responses of healthy subjects that experienced perturbations of their right upper limb introduced by a robotic manipulandum during a delayed-reach task. Our findings clarify the minimal preparatory time needed for goal-directed tuning of stretch reflex gains and reveal how these gains are modulated as a function of preparation duration.

## Methods

### Subjects

A total of 21 right-handed and neurologically healthy subjects (10 males of mean age 27.1 ±5.3 years and 11 females of mean age 24.1 ± 2.1 years) participated in the study. Data from four subjects were not included in statistical analyses, one due to poor adaptation to the task and three due to low-quality EMG signals. The remaining 17 subjects were eight males of mean age 27.6 ± 5.9 years and nine females of mean age 24.4 ± 2.0 years. All 21 subjects were financially compensated for their participation and gave informed, written consent before participating in the study. The subjects gave written informed consent, and the study was performed in line with the principles of the Declaration of Helsinki. It was approved by the Regional Ethical Review Board in Umeå.

### Experimental setup

Subjects were seated upright in a custom-designed adjustable chair positioned in front of the Kinarm End-Point Robot (BKIN Technologies). Using their right hand, they held onto the robotic manipulandum (Fig. 1a). The right forearm was supported within a tailored foam structure, which rested atop anan airsled system that permitted nearly frictionless movement within a two-dimensional workspace. To establish a stable mechanical linkage, the forearm, foam-mounted airsled, manipulandum handle, and hand were all secured together using a Velcro-fastened leather strap. This setup also ensured that the wrist remained aligned with the forearm in a neutral position throughout the experimental session. The robotic platform measured kinematic data regarding the hand’s position, and sensors inside the robotic handle recorded the forces exerted by the subjects’ right hand (six-axis force transducer; Mini40-R, ATI Industrial Automation). Position and force data were sampled at 1 kHz.

**Fig. 1.**
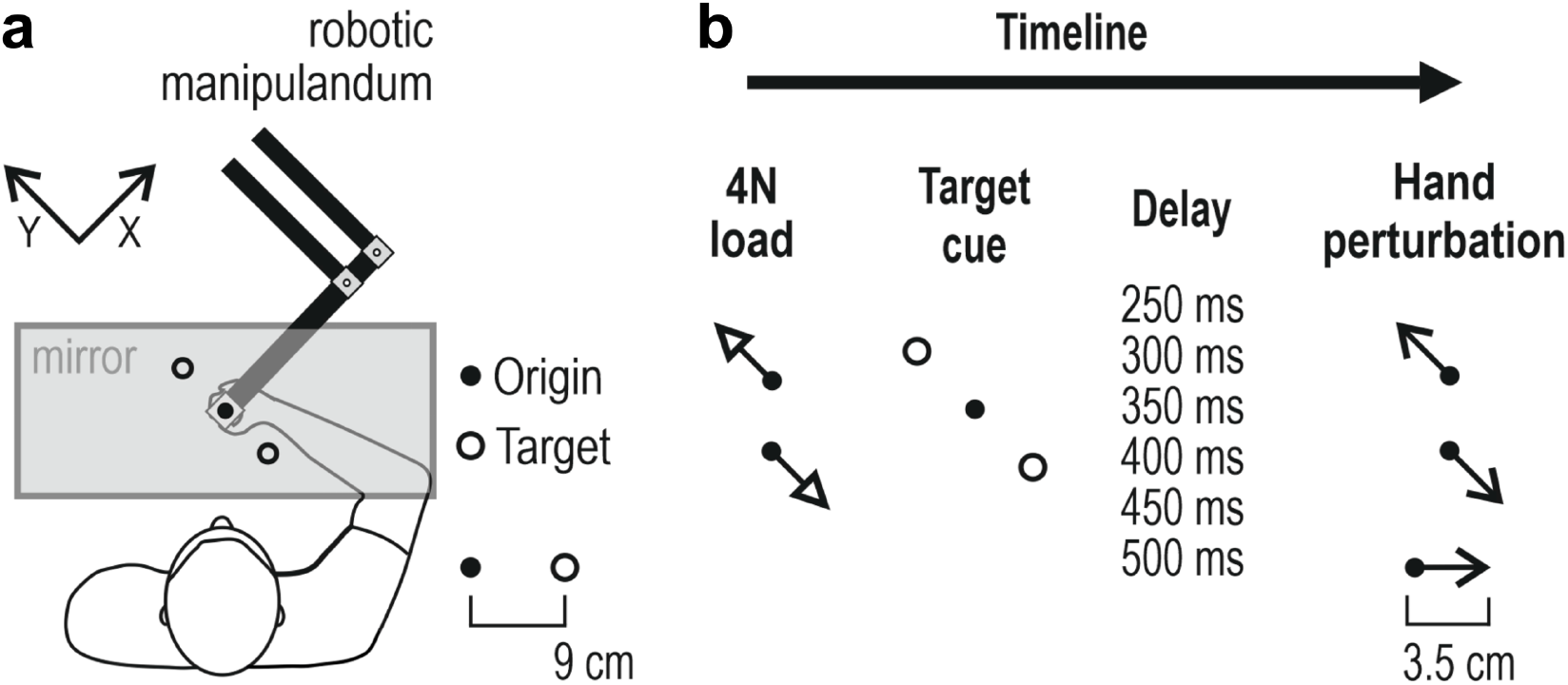
The robotic manipulandum and experimental timeline. (**a**) Subjects were seated upright in a custom-designed adjustable chair with their right dominant hand secured to a handle. The hand’s position was represented by a visual cursor representing where the origin and two visual targets were projected onto a mirror, preventing the subjects from seeing their hand (for details, see Methods). (**b**) Each trial began when the hand (i.e., cursor) was placed in the origin. A load (4N) was then applied in the front-and-left and right-and-back direction (135 and 315 degrees, respectively). During this phase, participants had to resist the load and maintain their hand at the origin. A visual target was then cued (through a red filled circle) for 250, 300, 350, 400, 450 or 500 ms. A position-controlled perturbation of the hand is then applied in the 135- or 315-degree direction (3.5 cm over 150 ms). The visual cursor’s position was still at the origin during the haptic perturbation. After the perturbation, the cued target turned into a green-filled circle corresponding to a “ Go” signal, and the subjects were instructed to move their hand such that the cursor hits the target

Bipolar surface EMG electrodes (Bagnoli DE-2.1, Delsys Inc., Natick, MA, USA) with 10.0×1.0 mm silver contacts and 10 mm inter-electrode distance were placed on the pectoralis major, posterior deltoid, and anterior deltoid muscles. Before placing the electrodes, the skin was cleaned with alcohol, and a double-sided adhesive tape was attached to the electrodes and coated with conductive gel. The electrodes were secured using surgical tape to ensure good contact throughout the session. A ground electrode (Dermatrode HE-R Reference Electrode type 00200–3400, 5 cm diameter, American Imex) was placed on the processus spinosus of the C7 region. The EMG signals were sampled at 1 kHz, bandpass filtered at 20-500 Hz, and A/D converted with 16-bit resolution using the Bagnoli EMG amplifier (Delsys Inc., Natick, MA, USA).

### Experimental design

The experimental design of the delayed-reach task involving haptic perturbations is summarised in Fig. 1 and is similar to what has been employed elsewhere, e.g., (Torell *et al*., 2023). Specifically, a one-way mirror prevented a view of the hand and robotic handle, and visual stimuli were projected in the plane of movement. Visual stimuli included a moving white cursor of 1 cm in diameter, representing the hand’s position. In addition, two targets (1.25 cm radius) located 9 cm from the centre point (‘origin’; 0.75 cm radius) were displayed as orange circle outlines unless cued. The targets were placed in a front-and-left and right-and-back direction, corresponding to 135 and 315 degrees.

Once the hand (i.e., the cursor) was placed in the origin, each trial started, where the cursor had to remain steady inside the origin for a randomly generated time (1 and 1.5 s), after which a 4N load with a 800 ms rise time and 1200 ms hold was applied in the 135 or 315 degree direction for 2 s. Then, one of the two targets turned into a filled red circle. After a pre-defined preparatory delay (250, 300, 350, 400, 450 or 500 ms), a position-controlled perturbation of the hand was applied in one of the two directions (3.5 cm with 150 ms rise-time and no hold). During the perturbation, the position of the cursor was fixed. At the onset of the perturbation, the filled red circle also turned green (the “ Go” signal). At the end of the perturbation, the subjects had to reach and remain within the target for 300 ms. After this, the subjects received visual feedback on their performance, and a correct trial was defined as reaching the target between 400 and 1400 ms after perturbation onset. Moving the cursor to the origin began the subsequent trial.

This study represents a full factorial design without centre points, i.e., each block contained 48 unique trials in a randomised order comprising two targets to be cued, two load directions, two perturbation directions, and six preparatory delays (2×2×2×6=48 trials). For the first five subjects, there was a total of 13 blocks, while for the latter subjects, there were 18 blocks, resulting in 864 trials. The subjects were instructed to take one-minute breaks after approximately consecutive blocks. However, the subjects were free to rest after completing any trial by pulling the cursor to the side of the workspace.

### Data processing

EMG data were high-pass filtered using a third-order, zero-phase-lag Butterworth filter with a 20 Hz cutoff and rectified. For each trial, the time between issuing the perturbation and movement onset (defined as 5% of peak hand velocity) was virtually identical across trials and subjects, given the nature of the hand perturbation (i.e., 18 ms). Therefore, all data were aligned on perturbation onset as determined by the KINARM and then shifted by a fixed delay of 18 ms. The raw EMG data were normalised by z-scoring, as described elsewhere (Dimitriou, 2014, 2016), to allow for inter-subject analysis. Briefly, this normalisation procedure involves concatenating all EMG signals from all blocks and calculating each muscle’s sample mean and standard deviation separately. We also removed the offset of the baseline around the perturbation (-25 to 25 ms) to ensure it was zero before averaging.

The first four blocks of trials were viewed as familiarisation trials and were not included in the subsequent analyses (see Results section). We removed trials (on average 7% across all participants) when the EMG or kinematics had a wiggly baseline when expected to be stable, e.g., due to the subject not being fully relaxed. For the EMG, a wiggly baseline within 200 and 0 ms before the perturbation was defined empirically as above 0.1 in mean absolute deviation. For the kinematics, the hand positions (both x- and y-axis) were defined empirically as outliers if they were 15 cm from the centre point within the first 15 ms after the perturbation.

Since this study focused on stretch reflex responses, we only analysed data from stretching muscles by analysing specific combinations of muscle and perturbation direction. To simplify analyses of individual muscles, the median EMG signal of each muscle was generated for each subject and trial type (i.e., for each load, perturbation, preparatory delay, and target direction), where the EMG signals were smoothed with a moving mean of five ms). Data directly representing goal-directed differences were generated by contrasting (subtracting) EMG values in trials where reaching the cued target required stretching the homonymous muscle vs. values from corresponding epochs in trials where reaching the target required muscle shortening.

### Statistical analysis

We analysed the goal-directed differences in stretch reflex responses for the different muscles and preparation delays using the Wilcoxon signed rank test with the False Discovery Rate (FDR) method to correct for multiple comparisons. We also tested the differences in the stretch reflex modulation between preparatory delays using the Kruskal-Wallis test with Fisher’s least significant difference (LSD) post-hoc tests. We used a significance level of 0.05. All data processing and statistics were performed using MATLAB (Version 2023b; MathWorks).

## Results

### Task performance

The subjects’ performance in the delayed-reach task reached a plateau relatively quickly, with the average performance in terms of % correct trials being high and stable after the first four blocks (Fig. 2a). Therefore, the first four blocks of trials were deemed familiarisation blocks and were not included in subsequent statistical analyses. Although the subjects generally performed well at the task, each subject occasionally initiated movements before receiving the ‘Go’ cue (2.6 ± 2.3 % of all trials), a phenomenon mainly observed during the longest preparatory delays, i.e., 450-500 ms (Fig. 2b). Although infrequent overall, these “ false starts” persisted despite repeated instructions to wait for the ‘Go’ cue. That is, the experimenter provided relevant instructions before data collection began and repeatedly after false starts, but interestingly, a relatively small number of such trials often persisted throughout the experiment.

**Fig. 2.**
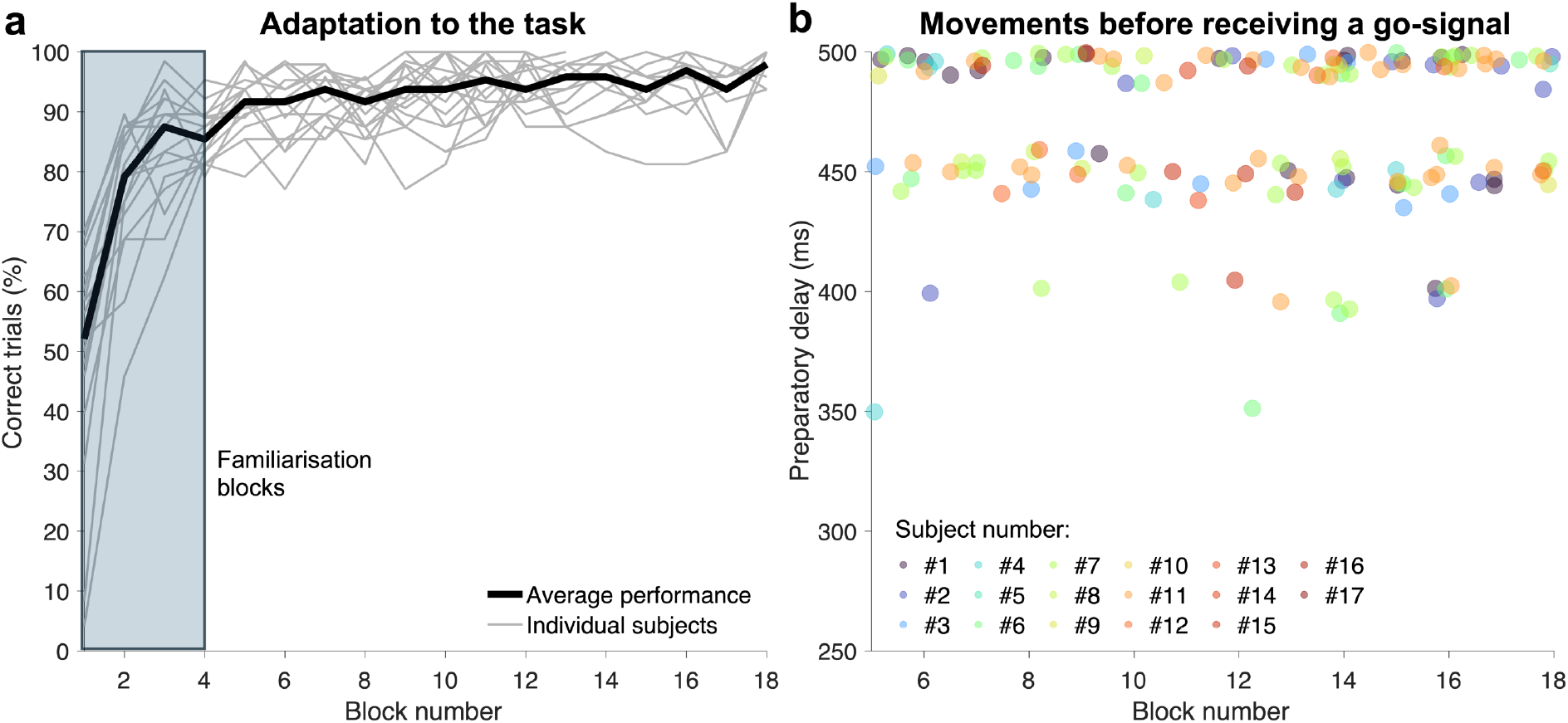
Reach performance and premature movement initiation. (**a**) The subjects’ performance at the task, in terms of % correct trials in each block of trials. A correct trial involved reaching the cued target within a certain time interval (see main text for more details). Performance reached a high and rather stable plateau after the first four blocks, which were deemed familiarisation blocks and not included in subsequent analyses. (**b**) Each subject occasionally initiated movements prematurely, i.e., before the ‘Go’ cue (2.6 ± 2.3 % of all trials), mainly during the longest preparatory delays (450-500 ms). These “ false starts” occurred even though the subjects were repeatedly instructed to wait for the ‘Go’ cue

### Goal-directed modulation of the stretch reflex responses – Pectoralis muscle

A representative example of hand positions, forces, and z-scored pectoralis EMG activity averaged across subjects for trials with a 400 ms preparatory delay is illustrated in Fig. 3a (unloaded pectoralis) and another in Fig. 3b (loaded pectoralis). As indicated in these figures, a preparatory delay of 400 ms is sufficient for inducing goal-directed tuning of SLR and LLR responses, at least when the homonymous muscle is unloaded (Fig. 3a). Also, note that the averaged hand positions and forces (along the x- and y-axis) are virtually identical during the haptic perturbation, regardless of the load direction (Fig. 3).

**Fig. 3.**
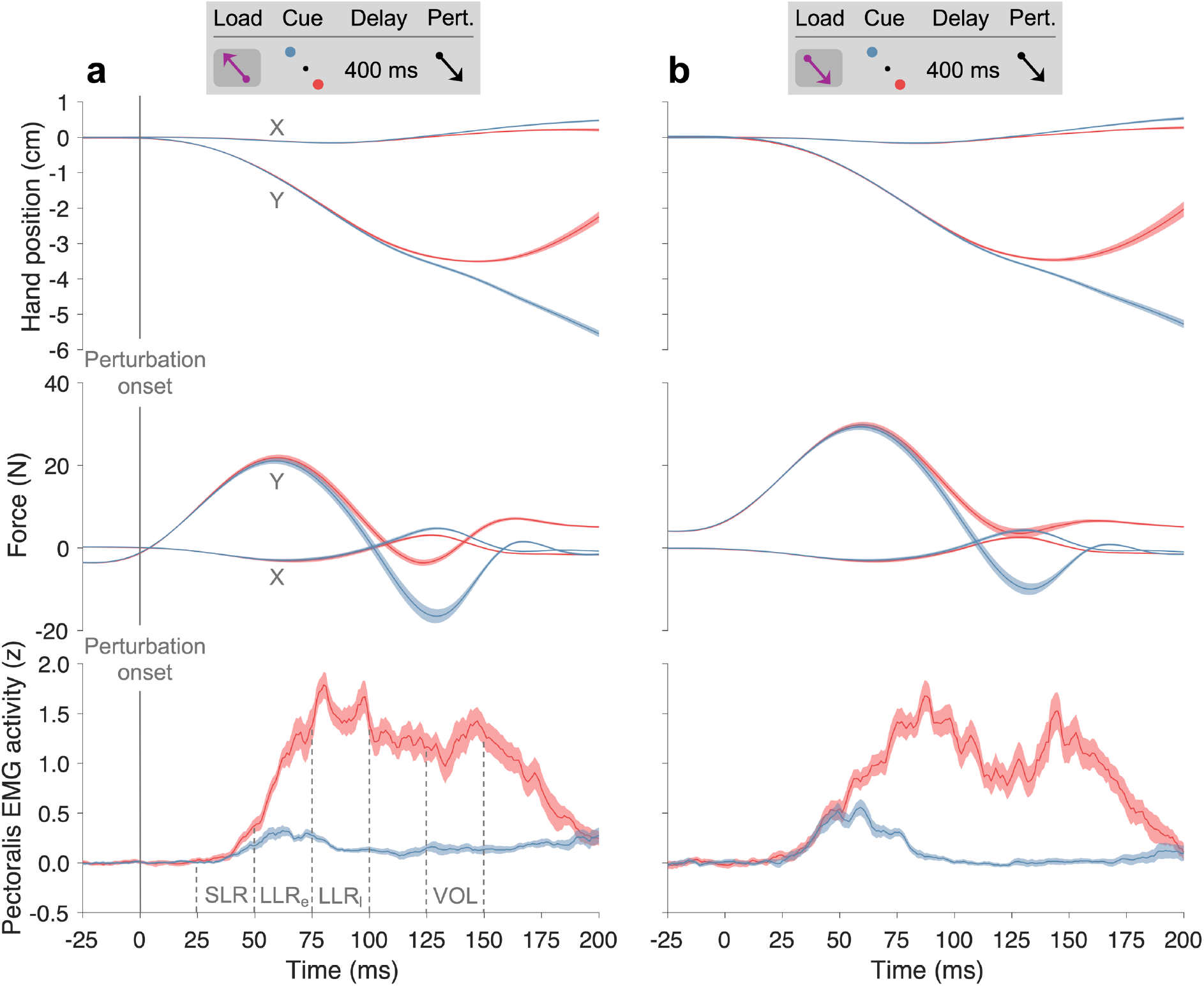
Averaged responses across subjects for trials with a preparatory delay of 400 ms. Averaged hand positions, haptic forces, and z-scored EMG activity across subjects for trials with a 400 ms preparatory delay when the pectoralis muscle was unloaded (**a**) and loaded (**b**). The vertical dashed lines indicate the epochs of the short- and long-latency stretch reflexes (i.e., SLR, early LLR, and late LLR) and the voluntary activity epoch (‘VOL’). Curve shading denotes ± 1 SE. In addition to z-transformation, the baseline offset of the EMG signals (pre-reflex activity) was removed at the level of single trials (see main text for more details)

Our analyses indicated that goal-directed pectoralis EMG activity across subjects varied as a function of preparatory delay (Fig. 4a). Analysis of individual EMG signals of the unloaded pectoralis (i.e., individual subject averages across trials) indicated a significant goal-directed difference in SLR (p < 0.05) for the 300-500 ms preparation delays (Fig. 4b). In contrast, we found no significant goal-directed difference in SLR when the preparatory delay was 250 ms. When the pectoralis was loaded, analyses indicated no goal-directed differences in SLR, regardless of the preparatory delay duration (Fig. 4c). Regarding the early and late LLR of the pectoralis muscle, analyses showed a significant goal-directed difference (p < 0.01) at all preparatory delays and load conditions (Fig. 4d-g).

**Fig. 4.**
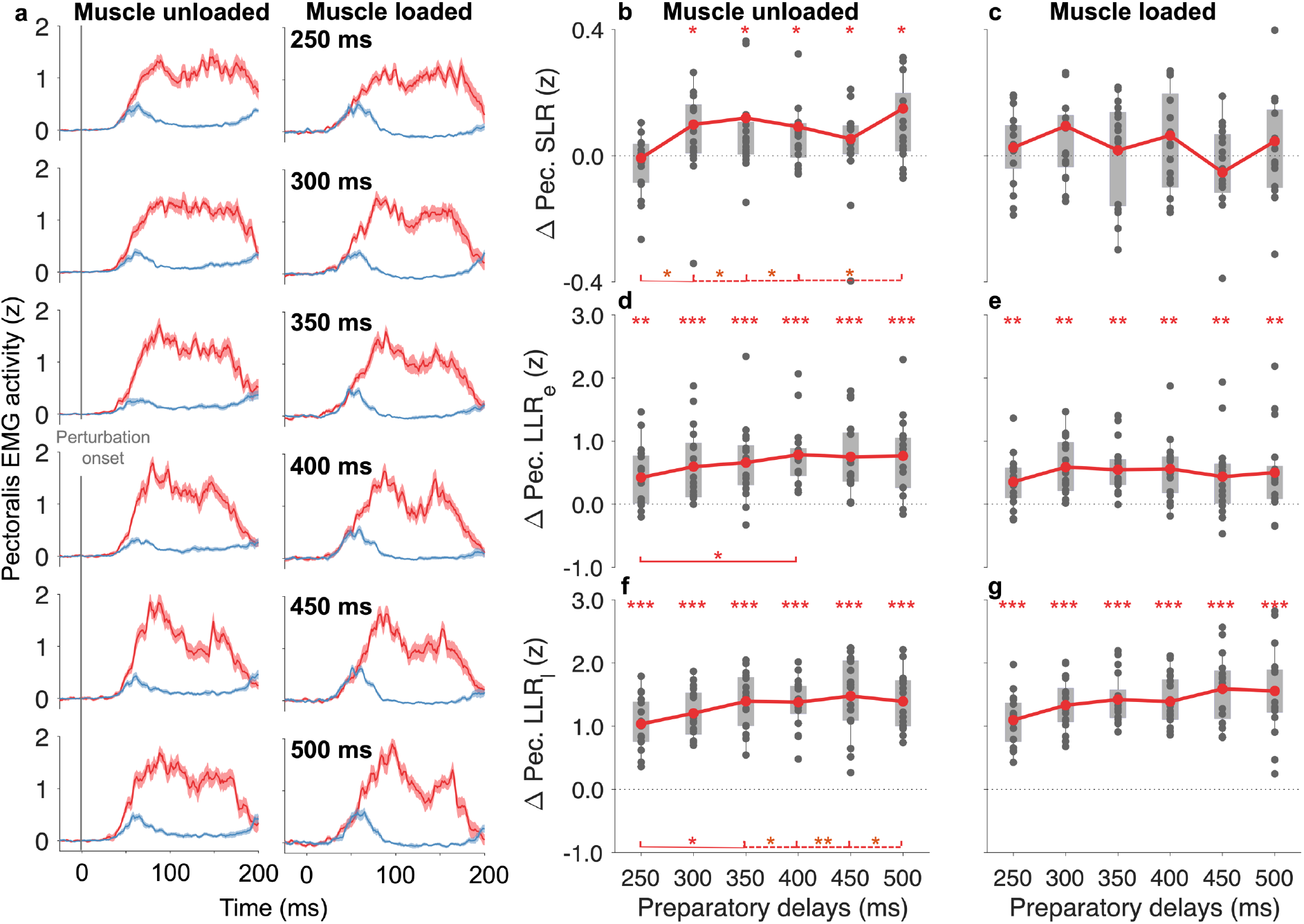
Goal-directed difference of pectoralis EMG activity. (**a**) Goal-directed difference of pectoralis EMG as a function of preparatory delay and load condition. (**b**) There was a goal-directed difference in SLR of the unloaded pectoralis following 300-500 ms preparatory delays. There was a significant difference in SLR EMG between the shortest preparatory delay (250 ms) and all other delays (p < 0.05), except for the 450 ms one (p = 0.057). (**c**) There were no goal-directed differences in SLR responses of the loaded pectoralis, regardless of preparatory delay duration. (**d-e**) There was a significant goal-directed difference in early LLR across all preparatory delays (p < 0.01) for both the unloaded and loaded muscles. However, there was only a significant difference in early LLR between the 250 and 400 ms delays (p < 0.05) when the muscle was unloaded. (**f-g**) There was a significant goal-directed difference in late LLR across all preparatory delays (p < 0.01) for both the unloaded and loaded muscles. There was a significant difference in late LLR between the shortest delay (250 ms) and all other delays (p < 0.05), except for the 300 ms (p = 0.31). Dots represent single-subject EMG values averaged across the relevant trials (see Methods), grey background rectangles represent upper and lower quartiles, and thin vertical lines represent 95% data range. * p < 0.05, ** p < 0.01, *** p < 0.001

Pairwise comparison of differences in SLR EMG in the unloaded pectoralis muscle as a function of preparation duration (Fig. 4b) indicated significantly weaker goal-directed tuning of SLR following the shortest delay (250 ms) vs. all other delays (p < 0.05), except for the 450 ms preparation delay (p = 0.057). For the early LLR (Fig. 4d), analyses revealed a significant difference in EMG activity only between the 250 and 400 ms delays (p < 0.05). For the late LLR (Fig. 4f), we found significantly weaker goal-directed tuning of EMG activity in the shortest preparatory delay vs. all other delays (p < 0.05), except for the 300 ms one (p = 0.31). Analyses indicated no significant difference in goal-directed tuning of voluntary EMG activity between the preparatory delays (not shown).

### Goal-directed modulation of the stretch reflex responses – Posterior and anterior deltoid muscles

There was a significant goal-directed difference in SLR EMG of the unloaded posterior deltoid muscle (i.e., pectoralis antagonist) following preparatory delays of 300-400 ms (p < 0.05; Fig. 5a). A pairwise comparison of differences in SLR EMG showed no significant difference between 300 and 400 ms preparatory delays (p > 0.05; Fig. 5a). In contrast, there was a significant difference between 350/400 ms and {250, 450, 500} ms (p < 0.05). For the early and late LLR, there was a significant goal-directed difference across all preparatory delays (p < 0.001; Fig. 5b-c). A pairwise comparison of differences in early LLR EMG showed a significant difference between the shortest delay (250 ms) and all other delays (p < 0.05), except for the 300 ms delay (p = 0.43). Analyses indicated no significant impact of preparation duration on the voluntary muscle activity (not shown).

**Fig. 5.**
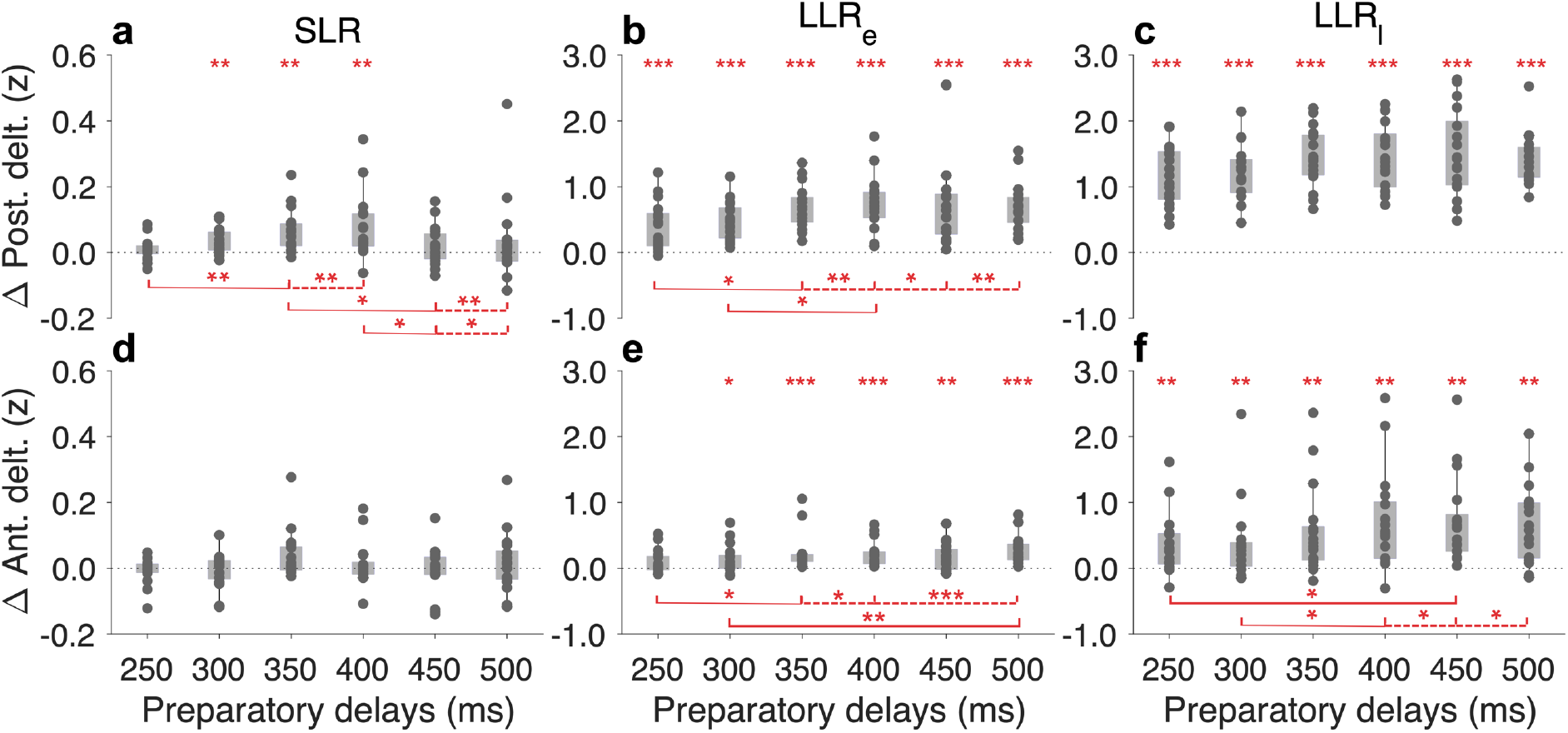
Goal-directed difference of the unloaded posterior and anterior deltoid muscles. (**a**) There was a goal-directed difference in SLR EMG of the posterior deltoid at preparatory delays of 300-400 ms (p < 0.05), and there was no significant difference in SLR EMG among those delays. In contrast, there was a significant difference between 350/400 ms and {250, 450, 500} ms (p < 0.05). (**b**-**c**) There was a goal-directed difference in early and late LLR EMG of the posterior deltoid across all preparatory delays (p < 0.001). For the early LLR activity, there was a significant difference between the shortest delay (250 ms) and all other delays (p < 0.05), except for the 300 ms delay (p = 0.43). (**d**) There was no goal-directed difference in the SLR of the anterior deltoid. (**e**-**f**) There was a goal-directed difference in early and late LLR EMG of the anterior deltoid across all preparatory delays (p < 0.001), except for the 250 ms delay in early LLR. For the early LLR (e), we found a significant difference between the shortest delay (250 ms) and all other delays (p < 0.05), except for 300 ms and 450 ms. For the late LLR (f), the longer preparatory delays were associated with a larger difference in EMG activity than the shorter delays. * p < 0.05, ** p < 0.01, *** p < 0.001

In contrast to the pectoralis and posterior deltoid muscles, there was no goal-directed difference in the SLR of the anterior deltoid (Fig. 5d). However, there was a goal-directed difference across all preparatory delays in the early and late LLR (p < 0.05; Fig. 5e-f), except for the 250 ms delay in early LLR (Fig. 5f). A pairwise comparison of differences in SLR EMG showed no significant difference between the preparatory delays (p > 0.05; Fig. 5d). For the early LLR (Fig. 5e), we found a significant difference between the shortest delay (250 ms) and all other delays (p < 0.05), except for 300 ms and 450 ms. For the late LLR (Fig. 5f), the longer preparatory delays were associated with a larger difference in EMG activity than the shorter delays. Finally, analyses indicated no significant impact of preparation duration on the voluntary muscle activity (not shown).

## Discussion

This study aimed to identify the minimum preparatory time required for goal-directed tuning of SLR and LLR responses and to characterise the incremental evolution of such tuning as a function of preparation duration. The two main findings of this study are that 1) a preparatory delay of 300 ms is sufficient for goal-directed tuning of SLR responses in the pectoralis and posterior deltoid muscles (e.g., Figs. 4-5) and 2) that an additional 100 ms (i.e., preparation of 400 ms) allows for even stronger goal-directed modulation of stretch reflex responses.

Our results confirm previous findings that preparatory delays of 250 ms are not associated with goal-directed SLR responses (Torell *et al*., 2023). While pertinent underlying mechanisms may already be at work during preparatory delays of 250 ms or shorter (Papaioannou and Dimitriou, 2021), longer preparatory delays are required for the systematic manifestation of goal-directed tuning of SLR responses, as assessed by surface EMG. Also, in line with previous findings, we show that unloading of the homonymous muscle facilitates the goal-directed tuning of SLRs. In contrast, muscle loading leads to ‘automatic’ gain-scaling of the SLR, regardless of goal and LLRs responses are robustly goal-dependent, regardless of background muscle loading (Pruszynski *et al*., 2009; Papaioannou and Dimitriou, 2021; Torell *et al*., 2023).

The optimal preparatory delay of 400 ms, found here, is approximately 200 ms longer than the goal-directed modulation of reflexes compared to what Yang and colleagues found (Yang *et al*., 2011). This difference is likely related to methodological differences and our participants preparing to perform a multi-joint task, not a single-joint task. Single-joint movements allow for a more controlled study of the preparatory processes. However, multi-joint movements provide a more naturalistic context for studying how the sensorimotor system coordinates complex actions, such as reach-to-grasp. In other words, it appears that the nervous system needs an additional ∼200 ms to sufficiently incorporate the goal-directed preparation of multi-joint reaching vs. single-joint movement.

Interestingly, for preparatory delays longer than 400 ms, the goal-directed tuning of the SLR (at least) seems to drop, and such longer preparatory delays are associated with more frequent “ false starts”, i.e., trials with premature movement initiation (i.e., before receiving a “ Go” signal). False starts occurred even though the subjects were instructed to wait for the “ Go” signal before and after these occurred, suggesting the engagement of specific cognitive mechanisms when the preparatory delay is long enough. The false start trials could occur for a myriad of reasons. When a series of alternative preparatory durations are involved in a task, given sufficient preparatory time the participants might explicitly or implicitly try to predict when the “ Go” cue will occur, and this process biasing their decision to initiate movement i.e., anticipating the Go cue to come sooner than it does for the longer preparatory delays. Some participants might tend to act more impulsively and, therefore, be more likely to act in anticipation of a “ Go” signal. When trying to wait, the CNS (and PNS) prepares for the upcoming movement. The cognitive preparation can sometimes “ leak” into premature muscle activation, resulting in a false start (Forgaard *et al*., 2019). Our findings suggest ≥ 450 ms preparatory delay is sufficient time for this cognitive motor leakage to occur in delayed reach.

In Figs. 4-5, we observed a decrease in goal-directed tuning of stretch reflex responses (Δ EMG activity) and voluntary activity, around 450 ms, for different preparation delays in the unloaded pectoralis muscle and posterior deltoid. The reason for this decrease is not directly apparent and requires further investigation. One can hypothesise that around 450 ms, there is a transient shift in the dominant preparatory processes. For instance, there might be an initial phase of more reflexive motor activity followed by a more specific phase of inhibition or refinement of the motor control policy involving higher cognitive functions. This could temporarily reduce the overall excitability of the reflex pathways. Preparatory delays longer than 450 ms could be sufficient for the higher cortical preparatory processes to dominate, and including EEG in future studies could shed light on this interesting phenomenon. On the other hand, having many different delays may engage a different mechanism than a task involving a discrete dichotomy in preparatory delays, i.e., long vs. short delay, with one being too short and one being sufficient. Indeed, this could explain why 250 ms was too little while 750 ms sufficed in a previous study (Torell *et al*., 2023), also characterised by a far smaller number of false-start trials.

In contrast to the case of the pectoralis and posterior deltoid muscle, there was no goal-directed tuning of SLR responses in the anterior deltoid muscle. In our task, the anterior deltoid functioned as a pectoralis synergist. This, in turn, may imply that, at least after task familiarisation, the sensorimotor system does not apply independent fusimotor control to all synergists potentially engaged in a task but rather to a subset of these. In other words, independent fusimotor control appears highly selective across the muscles that could be similarly engaged as agonists and antagonists in a motor task. Congruent with the idea that feedback controllers are loaded before movement initiation (Ahmadi-Pajouh, Towhidkhah and Shadmehr, 2012), we identify the minimum amount of time required by these controllers for manifesting goal-directed modulation of reflex responses, highlighting that feedback controllers likely also engage top-down fusimotor control (Dimitriou, 2022).

In the current study, reflex responses appear strongly inhibited or even absent under certain experimental conditions. This can be attributed to the context-dependent modulation of the stretch reflex. In some situations, the CNS might actively suppress or gate the stretch reflex to allow for controlled movement (Shemmell, An and Perreault, 2009). If muscle stretch occurs during such higher-order suppression or gating, the resulting contraction might be weaker than expected, potentially being perceived as absent or a minimal response. In the current study, the task strives to be as naturalistic as possible, allowing the subjects to perform the task without being too constrained, which may result in a different recruitment of motor units (Hodson-Tole and Wakeling, 2009). Since we used bipolar surface EMG electrodes to quantify the stretch reflex responses in this naturalistic task, this may partly explain the variability in reflex responses across participants. To overcome this limitation, using multi-channel surface EMG to cover a larger part of the muscle would lead to higher signal-to-noise estimates of the stretch reflex responses.

In conclusion, we examined how multiple preparatory delays impact the goal-directed modulation of stretch reflex gains of the pectoralis, anterior and posterior deltoid muscles in a multi-joint reaching task. We found that a preparatory delay of 300 ms is sufficient for goal-directed tuning of SLR responses in the pectoralis and posterior deltoid muscles, and 400 ms allows for even stronger goal-directed modulation of stretch reflex responses. Our results clarify the minimum preparatory time required for goal-directed tuning of stretch reflex gains at the level of the dominant upper limb and characterise the relationship between reflex gains and preparation duration. We show that this relationship is not strictly linear, likely reflecting the interplay of multiple feedback mechanisms functioning at different time frames.

## Acknowledgements

We thank Carola Hjältén and Anders Bäckström for their technical support. This work was supported by grants awarded to M.D. by Hjärnfonden (FO2024-0425-HK-88) and R.R. by the Swedish Research Council for Sport Science (P2025-0173).

## Declarations

### Consent to participate and publish

The subjects gave written informed consent to participate in this study. The authors affirm that human research participants provided informed consent to submit the manuscript to a scientific journal.

### Ethics approval

The subjects gave written informed consent, and the study was performed in line with the principles of the Declaration of Helsinki. It was approved by the Regional Ethical Review Board in Umeå.

### Data and code availability

Data and source code are available upon request.

### Authors’ contribution statements

All authors contributed to the study’s conception and design. RR performed data curation and analysis. RR wrote the first draft of the manuscript. All authors reviewed and edited the manuscript. All authors read and approved the final manuscript.

